# Three-dimensional understanding of the morphological complexity of the human uterine endometrium

**DOI:** 10.1101/2020.05.29.118034

**Authors:** Manako Yamaguchi, Kosuke Yoshihara, Kazuaki Suda, Hirofumi Nakaoka, Nozomi Yachida, Haruka Ueda, Kentaro Sugino, Yutaro Mori, Kaoru Yamawaki, Ryo Tamura, Tatsuya Ishiguro, Teiichi Motoyama, Yu Watanabe, Shujiro Okuda, Kazuki Tainaka, Takayuki Enomoto

**Author notes:** Correspondence (KY), (KT) and (TE).

## Abstract

The histological basis of the human uterine endometrium has been established by 2D observation. However, the fundamental morphology of endometrial glands is not sufficiently understood because these glands have complicated winding and branching patterns. To construct a big picture of endometrial gland structure, we performed tissue-clearing-based 3D imaging of human uterine endometrial tissue. Our 3D immunohistochemistry and 3D layer analyses revealed that endometrial glands formed a plexus network in the stratum basalis, similar to the rhizome of grass. We then extended our method to assess the 3D morphology of adenomyosis, a representative “endometrium-related disease”, and observed 3D morphological features including direct invasion of endometrial glands into the myometrium and an ant colony-like network of ectopic endometrial glands within the myometrium. Thus, 3D analysis of the human endometrium and endometrium-related diseases will be a promising approach to better understand the pathologic physiology of the human endometrium.

## Introduction

The human endometrium is a dynamic tissue that exhibits high regenerability after cyclic shedding, namely, menstruation. Menstruation is a unique biological phenomenon that occurs in a limited number of mammals, such as humans and other higher primates (Emera et al., 2012; Garry et al., 2009). Menstruation involves cyclic morphological and functional changes in the uterine endometrium that occur on a monthly basis in response to ovarian hormones (Noyes et al., 1950). The uterine endometrium changes dramatically based on the phases of the menstrual cycle (i.e., the proliferative phase, the secretory phase, and menstruation) and plays a crucial role in the implantation of fertilized eggs. On the other hand, “endometrium-related diseases” such as adenomyosis, endometriosis, endometrial hyperplasia, and endometrial cancer originate in the uterine endometrium due to its high intrinsic regenerative capacity, and these diseases affect the lives of women from puberty until after menopause (Garcia-Solares et al., 2018; Koninckx et al., 2019; Morice et al., 2016). However, the pathogenesis of endometrium-related diseases remains unclear, and further investigations focusing on the endometrium from the standpoint of disease prevention are required.

The conventional morphological theory of endometrial structure has been established based on two-dimensional (2D) histological observation (Johannisson et al., 1982; McLennan and Rydell, 1965; Noyes et al., 1950). Histologically, the endometrium is lined by a simple luminal epithelium and contains tubular glands that radiate through endometrial stroma toward the myometrium via coiling and branching morphogenesis (Gray et al., 2001). The human endometrium is stratified into two zones: the stratum functionalis and the stratum basalis. The stratum functionalis is shed during menstruation and regenerates from an underlying basal layer during the proliferative period. Therefore, it is widely assumed that regeneration of the stratum functionalis depends on endometrial progenitor/stem cells residing in the stratum basalis (Kyo et al., 2011; Maruyama and Yoshimura, 2012; Padykula, 1991; Prianishnikov, 1978). Despite this well-established understanding, neither the detailed mechanisms of endometrial regeneration during the menstrual cycle nor the localization of endometrial progenitor/stem cells have been fully characterized (Garry et al., 2010; Gellersen and Brosens, 2014; Santamaria et al., 2018). One of the reasons for this knowledge gap is that the fundamental structure of the human endometrium has not been sufficiently clarified. Since human endometrial glands have complicated winding and branching morphologies, it is extremely difficult to assess the whole shapes of glands by only 2D histopathology imaging.

In our previous genomic study, sequence analysis of 109 single endometrial glands revealed that each gland carried distinct somatic mutations in cancer-associated genes such as *PIK3CA*, *KRAS*, and *PTEN* (Suda et al., 2018). Remarkably, the high mutant allele frequencies of somatic mutations per endometrial gland have indicated the monoclonality of each gland. The presence of cancer-associated gene mutations in histologically normal endometrial glands provides important clues on the pathogenesis of endometrium-related diseases. Hence, we hypothesized that clonal genomic alterations in histologically normal endometrial glands may change the stereoscopic structure of the uterine endometrium, leading to susceptibility to endometrium-related diseases. To this end, we initially needed to evaluate and understand the three-dimensional (3D) morphology of the normal uterine endometrium.

Recently, several tissue-clearing methods have been developed to enable 3D imaging of rodent and primate tissue samples (Chung et al., 2013; Erturk et al., 2012; Hama et al., 2011; Ke et al., 2013; Susaki et al., 2015; Tainaka et al., 2016). The combination of these methods with the use of various types of optical microscopy including confocal fluorescence microscopy and light-sheet fluorescence (LSF) microscopy enables us to reconstitute 3D views of the tissues and provide 2D images of free-angle sections without the process of tissue slicing. In this study, we applied the updated clear, unobstructed brain/body imaging cocktails and computational analysis (CUBIC) protocol with fluorescent staining of clear full-thickness human uterine endometrial tissue because CUBIC has several advantages that make it suitable for our analysis (Tainaka et al., 2018). First, CUBIC is excellent in terms of its ease of use and safety because CUBIC is a hydrophilic tissue-clearing method. Second, the updated CUBIC protocols offer fast and effective clearing of various human tissue (Tainaka et al., 2018). Therefore, we determined that CUBIC had proper capabilities for clearing human uterine tissue.

Here, we aimed to establish a novel method for evaluating the 3D structure of the uterine endometrium and to define the “normal” 3D morphology of the endometrial glands. We successfully obtained 3D full-thickness images of the human uterine endometrium by combining an updated CUBIC protocol with the use of LFS microscopy. We also succeeded in constructing the 3D morphology of the glands and developed the new concept of 2D-shape images of the human endometrium. Finally, we used our 3D imaging method to reveal the 3D pathologic morphologies of adenomyosis, a representative “endometrium-related diseases”. Elucidation of the 3D structure of the human endometrial glands will provide further insights into various fields, including histology, pathology, pathophysiology and oncology.

## Results

### Tissue clearing and 3D imaging of human uterine tissue by updated CUBIC protocol

To clear human uterine endometrial tissue, we applied CUBIC protocol IV, which was previously utilized for clearing human brain tissue (Figure 1A) (Tainaka et al., 2018). We collected 14 uterine endometrial samples from 12 patients who underwent hysterectomy due to gynecological diseases with no lesions in the endometrium (Figure 1B, Table 1). With CUBIC protocol IV, we succeeded in substantially clearing all 14 human uterine endometrial tissues (Figure 1C). The autofluorescence signal derived from collagen and elastic fibers (Zhao et al., 2020) was useful for observing the intact 3D structure of uterine endometrial tissue by LSF microscopy (Figure 1D left panel). We added immunostaining with a fluorescently labeled anti-cytokeratin (CK) 7 antibody to highlight the endometrial gland structure. Immunohistochemical staining with the anti-CK7 antibody demonstrated selective labeling of the luminal and glandular epithelial cells running through the endometrial stroma with single-cell resolution (Figure 1D and Figure S1). By 3D reconstitution of the LSF microscopic images of the CK7-stained human endometrium, we succeeded in visualizing detailed 3D structures of endometrial glands (Figure 1E). Our stereoscopic image made it possible to analyze free-angle images of cross-sections. As shown in Figure 1F, the XY slice of uterine endometrial tissue after implementation of the CUBIC protocol retained the characteristic 2D morphology of endometrial glands for each phase, namely, curving glands in the proliferative phase and serrated glands in the secretory phase. The 3D image reconstituted by Imaris software (Bitplane) enabled us to observe continuous tomographic images of the human endometrium in all directions (Video S1).

**Figure 1.**
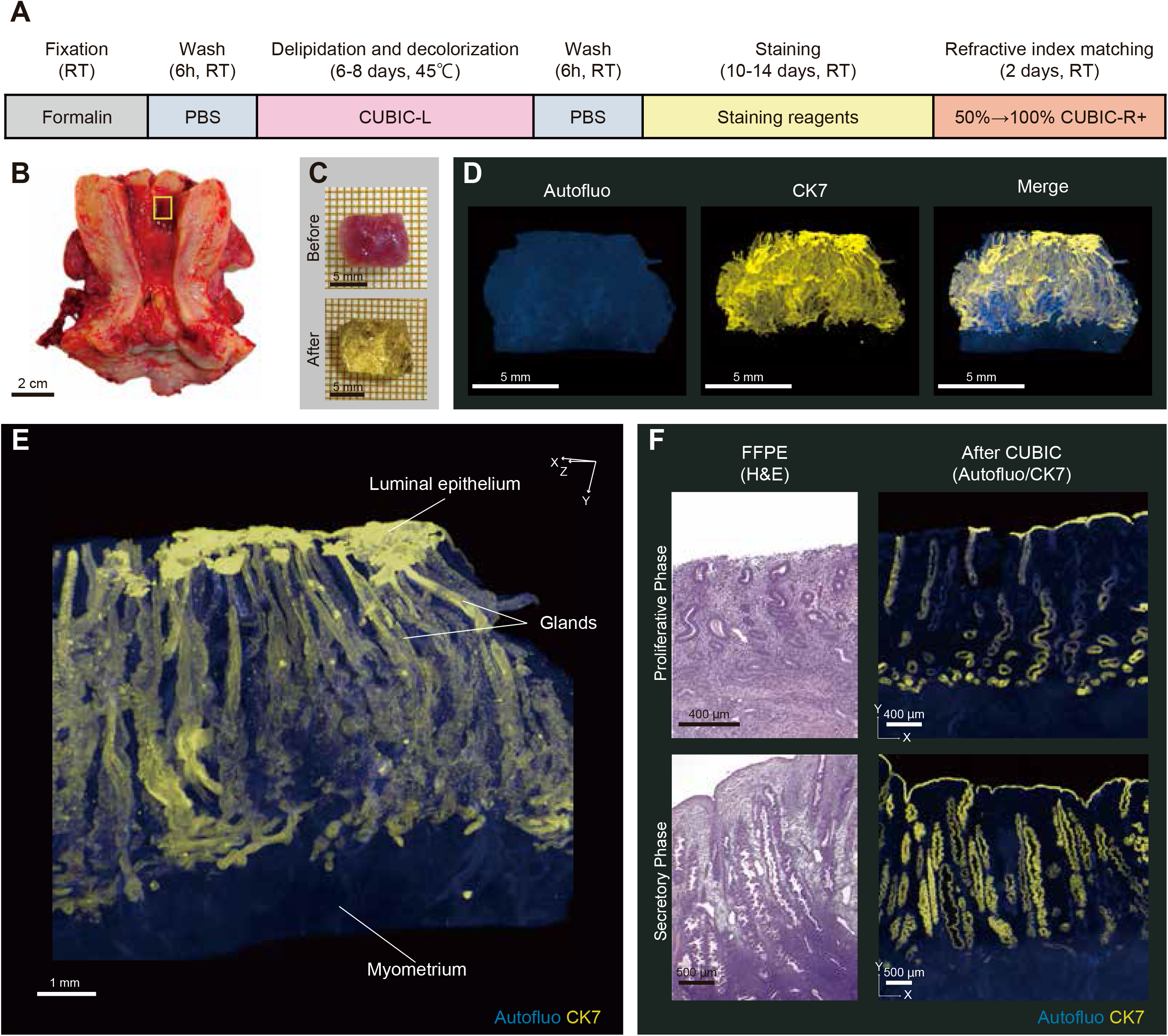
Tissue cleaning and 3D imaging of human uterine tissue using CUBIC. (A) Schematic diagram of the clearing and immunostaining protocol for human uterine tissue. (B) Sampling site (yellow box) of human uterine tissue from subject E2. (C) Cleaning performance of CUBIC protocol □ for human uterine tissue from subject E2. (D) 3D images of subject E2 stained with Alexa Fluor 555-conjugated anti-CK7 antibody with clearing by CUBIC. (E) Magnified 3D distribution of subject E2. (F) Comparison of a microscopic image of FFPE tissue after H&E staining and the reconstituted XY-section image after clearing by CUBIC. FFPE made by adjacent uterine tissue used for whole-mount 3D analysis. Upper panels: images of endometrium in proliferative phase (subject E1). Lower panels: images of endometrium in the secretory phase (subject E8). XY plane optical slices (subject E1, z = 7.62 μm; subject E8, z = 6.61 μm). (D-F) Images were obtained by LSF microscopy. Autofluorescence was measured by excitation at 488 nm. The CK7-expressing endometrial epithelial cells were measured by excitation at 532 nm lasers. RT, room temperature; Autofluo, autofluorescence; CK7, cytokeratin 7; FFPE, formalin-fixed paraffin-embedded; H&E, hematoxylin and eosin See also Figure S1.

**Table 1.** Clinical characteristics of subjects.

| Subject number | Age | Clinical diagnosis | Menstrual cycle | Gravidity | Parity | BMI |
| --- | --- | --- | --- | --- | --- | --- |
| E1 | 30 | Cervical cancer ( I A1) | Proliferative phase | 1 | 1 | 34.1 |
| E2 | 39 | Cervical cancer ( I B1) | Proliferative phase | 3 | 2 | 17.9 |
| E3 | 43 | Pelvic organ prolapse | Proliferative phase | 1 | 1 | 23.8 |
| E4 | 46 | Myoma uteri | Proliferative phase | 0 | 0 | 25.0 |
| E5-1, 2 | 42 | Myoma uteri | Secretory phase | 4 | 2 | 20.7 |
| E6 | 44 | Myoma uteri, pelvic organ prolapse | Secretory phase | 1 | 1 | 22.8 |
| E7 | 44 | Myoma uteri | Secretory phase | 0 | 0 | 29.9 |
| E8 | 46 | Malignant transformation within a dermoid cyst | Secretory phase | 5 | 4 | 22.5 |
| E9 | 46 | Cervical cancer ( I B1) | Secretory phase | 1 | 1 | 17.8 |
| E10 | 49 | Myoma uteri, pelvic organ prolapse | Secretory phase | 3 | 3 | 20.9 |
| E11-1, 2 | 45 | Myoma uteri | Menstrual phase | 3 | 3 | 28.7 |
| E12 | 48 | Myoma uteri | Menstrual phase | 0 | 0 | 25.2 |
| A1 | 42 | Adenomyosis | Secretory phase | 2 | 0 | 21.2 |
| A2 | 45 | Adenomyosis | Secretory phase | 0 | 0 | 23.8 |
| A3 | 42 | Adenomyosis | undergoing GnRH agonist treatment | 2 | 1 | 26.4 |

### Human endometrial glands are composed of occluded glands and plexus of glands

Our 3D imaging of human uterine endometrial tissue clarified some unique 3D morphologies of endometrial glands that had not been detected by 2D histological observation. First, we detected an occluded gland by observation of continuous tomographic images (Figure 2A). To visualize and assess the 3D morphology of the occluded gland, we added pseudocolor to the gland in the 3D image independently (Figure 2B, see the STAR methods for details). Intriguingly, the occluded gland rose from the bottom of the endometrium together with other nonoccluded glands but swelled up without reaching the luminal epithelium (Figure 2C).

**Figure 2.**
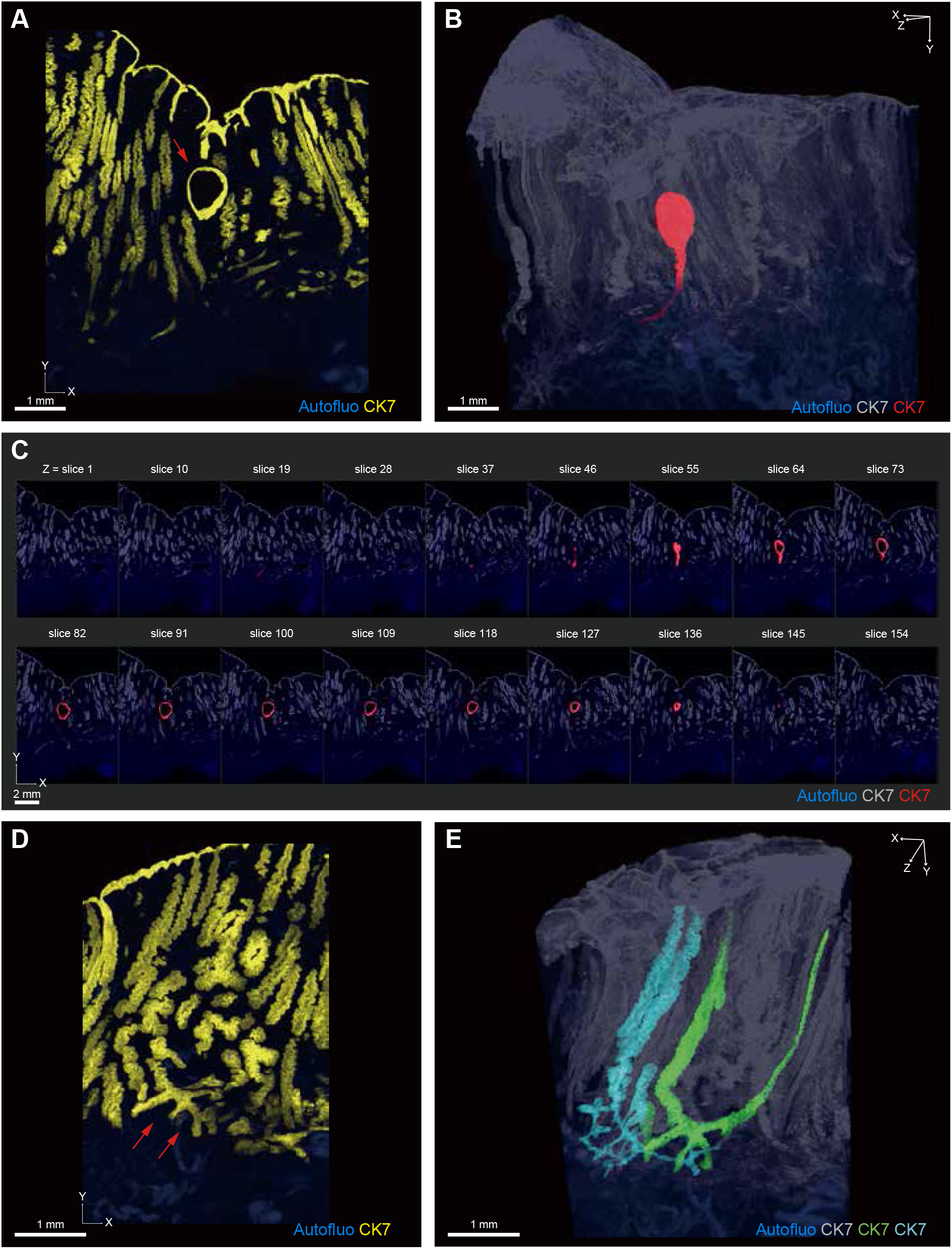
Characteristic 3D morphology of human endometrial glands. (A-C) The occulated gland (subject E8). (A) The reconstructed XY-section images (z = 99 μm). The red arrow indicates an occulated gland. (B) The reconstituted 3D distribution of the occulated gland that was pseudocolored and separated as a new channel by the Surface module in the Imaris software. (C) The reconstituted XY-section images (z-stack: 6.61 μm/slice) for every 9 slices of the reconstituted image shown in panel B. (D, E) The branches of endometrial glands (subject E8). (D) The reconstituted XY-section images (z = 198 μm). Red arrows indicate the branches. (E) The reconstituted 3D distribution of the branched glands that were pseudocolored and separated as new channels by the Surface module in the Imaris software. Images were obtained by LSF microscopy. Autofluorescence was measured by excitation at 488 nm. The CK7-expressing endometrial epithelial cells were measured by excitation at 532 nm lasers. Autofluo, autofluorescence; CK7, cytokeratin 7 See also Figures S2 and S3.

Second, many branches of endometrial glands were detected at the bottom of the endometrium (Figure 2D). Until now, it has been widely assumed that human endometrial glands branch from a single gland and radiate through endometrial stroma toward the myometrium on the basis of previous 2D histological studies (Cooke et al., 2013; Gray et al., 2001). However, our 3D imaging revealed a more complex network of endometrial glands. When we assigned pseudocolors to two endometrial glands lying next to each other by Imaris software, we uncovered a plexus of endometrial glands near the bottom of the endometrium. Some endometrial glands shared the plexus and rose toward the luminal epithelium (Figure 2E, Video S2). The occluded glands and the plexus of glands were observed in all samples of the proliferative phase (n = 4) and secretory phase (n = 7) (Figure S2, S3), suggesting that they were basic components of the normal human endometrium.

### Plexus structure of the glands is mainly located in the lower layers

According to the 3D morphology observation, plexuses of the glands seemed to exist at the lower part of the endometrium. To shed light on where many plexuses of the glands were formed, it was necessary to divide an endometrium into layers and assess the structure of endometrial glands per layer. Since the boundary between the endometrium and the myometrium was not flat, linear XZ sections were unsuitable to evaluate the 3D-layer distribution of endometrial glands. Therefore, we manually traced the borderline between the endometrium and myometrium of the XY section and made a reconstituted 3D bottom surface of the endometrium (Figure 3A left and middle panels, Figure S4A-B, see the STAR methods for details). Then, new 3D layers of endometrial glands were created to be the same distance from the bottom layer and have a thickness of 150 μm (Figure 3A right panel, Figure S4C-F). We extracted five 3D layers from the bottom to the lumen of the endometrium (layer 1: 1-150 μm, layer 2: 151-300 μm, layer 3: 501-650 μm, layer 4: 1001-1150 μm, layer 5: 1501-1650 μm). The plexus structure of the glands was mainly located in the lower layers (layers 1 and 2), and the morphology of the glands became columnar as the layers approached the lumen of the endometrium (layers 3-5).

**Figure 3.**
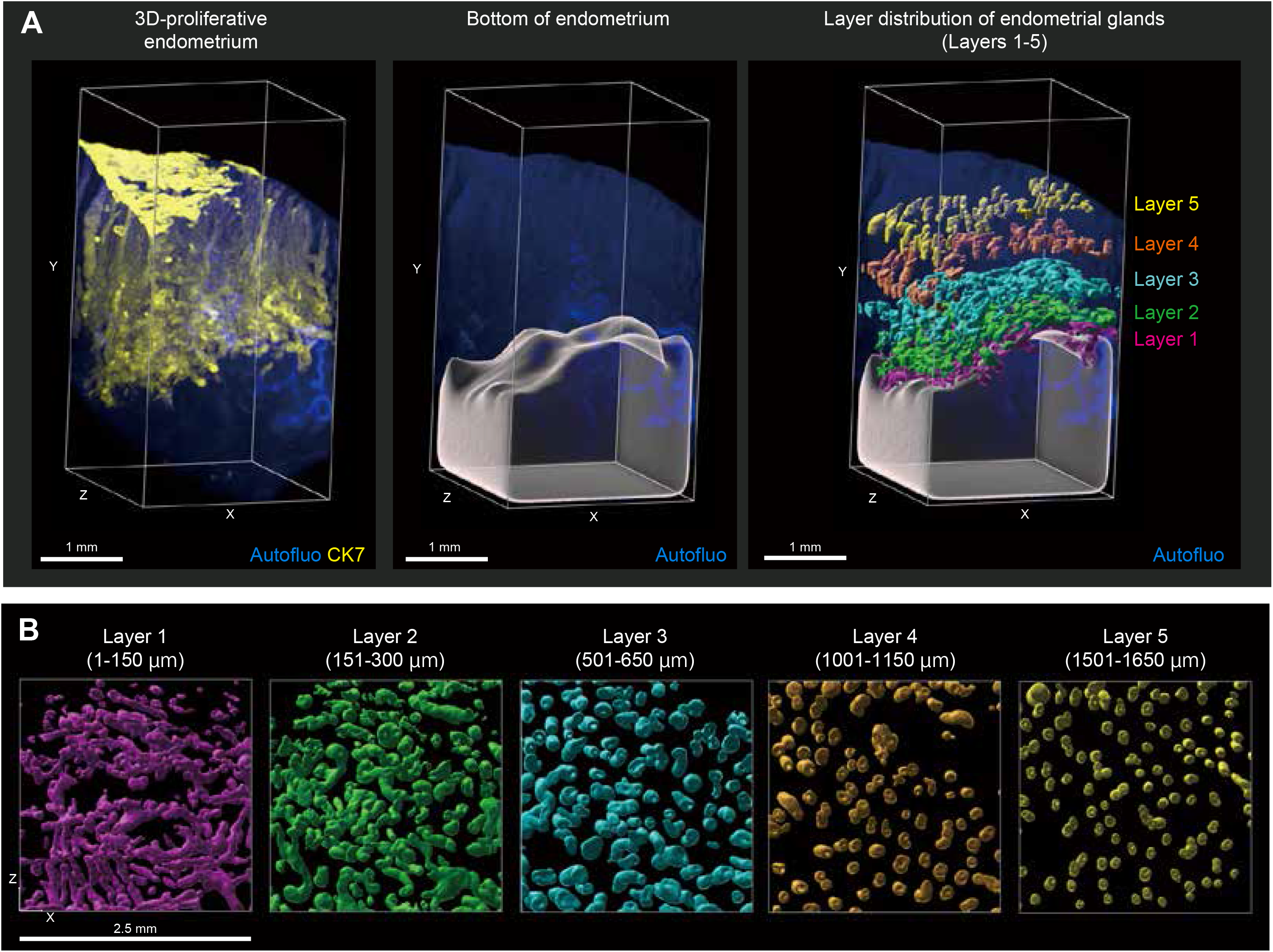
3D-layer distribution of human endometrial glands. (A) Left panel: the 3D tissue image was cropped in the XZ plane to 2.5 × 2.5 mm (subject E1). Middle panel: The reconstituted 3D bottom layer of the endometrium. Right panel: 3D layers of endometrial glands were created at the same distance from the bottom layer and with a thickness of 150 μm by the Surface module in the Imaris software. Layer 1 (magenta): 1-150 μm; layer 2 (green): 151-300 μm; layer 3 (light blue): 501-650 μm; layer 4 (orange): 1001-1150 μm; and layer 5 (yellow): 1501-1650 μm. (B) The XZ plane view (y = 150 μm) of five layers made by the Surface module in the Imaris software. After surface extraction, each structure was manually curated, and extra surface signals were eliminated. Images were obtained by LSF microscopy. Autofluorescence was measured by excitation at 488 nm. The CK7-expressing endometrial epithelial cells were measured by excitation at 532 nm lasers. Autofluo, autofluorescence; CK7, cytokeratin 7 See also figure S4.

### Plexus structure in the stratum basalis is preserved in menstrual phase

In menstruation, the stratum functionalis exfoliates, and the stratum basalis remains. To uncover whether the plexus of endometrial glands was located in the stratum basalis, we observed a 3D-layer distribution of endometrial glands in menstruation (menstrual cycle day 2) (Figure 4A, B, C). We reconstituted the bottom surface of the endometrium in a 3D image and divided the endometrium into two 3D layers (layer 1: 1-150 μm, layer 2: 151-300 μm). In menstruation, the plexus structure of the glands remained at the bottom of the endometrium (Figure 4D, Video S3). The other two samples obtained during menstruation also had similar plexus structures in their lower layers (Figure S5). These results revealed that the plexus structure of endometrial glands is mainly located in the stratum basalis regardless of menstrual cycle phase.

**Figure 4.**
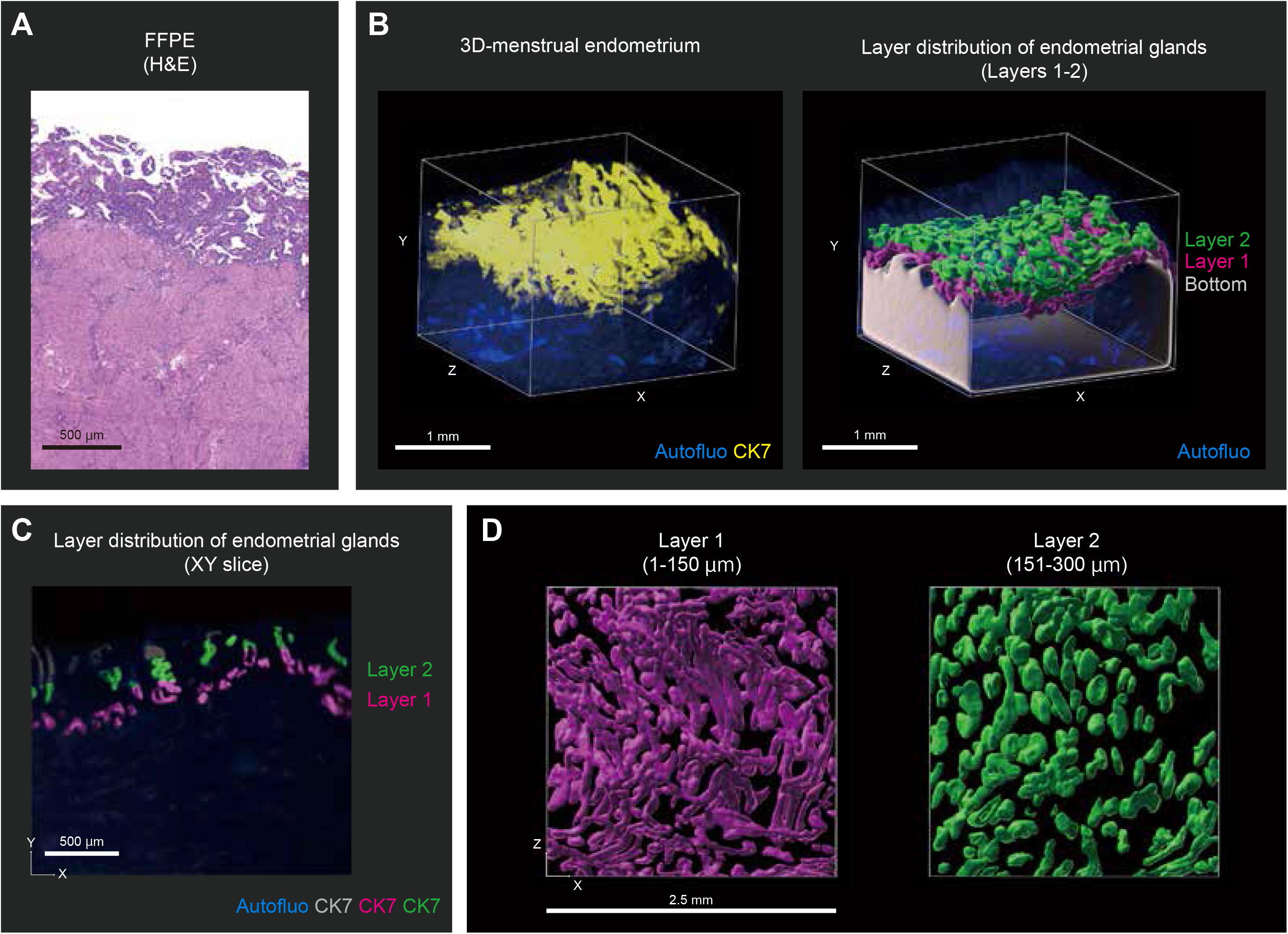
3D-layer distribution of endometrial glands in a menstruation case. (A) The microscopic image of FFPE tissue after H&E staining of the menstruation case (subject E11-1). (B) Left panel: the 3D tissue image was cropped in the XZ plane to 2.5 × 2.5 mm (subject E1-1). Right panel: The reconstituted 3D bottom layer of the endometrium and 3D layers of endometrial glands were created at the same distance from the bottom layer and with a thickness of 150 μm by the Surface module in the Imaris software. Layer 1 (magenta): 1-150 μm; layer 2 (green): 151-300 μm. (C) The reconstituted XY-section images (z = 47.6 μm) and each layer were pseudocolored and separated as new channels by the Surface module in the Imaris software. (D) The XZ plane view (y = 150 μm) of two layers made by the Surface module in the Imaris software. After surface extraction, each structure was manually curated, and extra surface signals were eliminated. Images were obtained by LSF microscopy. Autofluorescence was measured by excitation at 488 nm. The CK7-expressing endometrial epithelial cells were measured by excitation at 532 nm lasers. FFPE, formalin-fixed paraffin-embedded; H&E, hematoxylin and eosin; Autofluo, autofluorescence; CK7, cytokeratin 7 See also figure S5.

### Adenomyosis is stereoscopically characterized by ant colony-like network and direct invasion of endometrial glands into the myometrium

Finally, we applied our method for 3D visualization of endometrial glands to visualizing tissues in the context of adenomyosis, a benign gynecologic disease that is characterized by the presence of ectopic endometrial tissue within the myometrium. We collected adenomyosis tissue samples from three patients who underwent hysterectomy (Table 1). Two subjects (A1 and A2) did not receive any hormonal therapy within three years before the operation. Subject A3 was under therapy with a gonadotropin-releasing hormone (GnRH) agonist. With the application of CUBIC protocol IV (Figure 1A), the adenomyosis tissue samples were successfully cleared and stained with the anti-CK7 antibody (Figure 5A, Figure S6). The reconstituted 3D image of adenomyosis showed the detailed 3D structures of ectopic endometrial tissues within the myometrium. Interestingly, the ectopic endometrial glands had lengthened thin branches and an expanded adenomyotic lesion. As a result, the adenomyotic lesion that formed was similar to an ant colony within the myometrium (Figure 5B and 5C). This ant colony-like structure was more remarkable in subjects A1 and A2 than in subject A3, who had undergone GnRH agonist therapy (Figure 5D). Furthermore, we observed the direct invasion of the eutopic endometrial gland into the myometrium, leading to the formation of adenomyotic lesions in subjects A1 and A3 regardless of GnRH agonist therapy (Figure 5B and 5D, Video S4).

**Figure 5.**
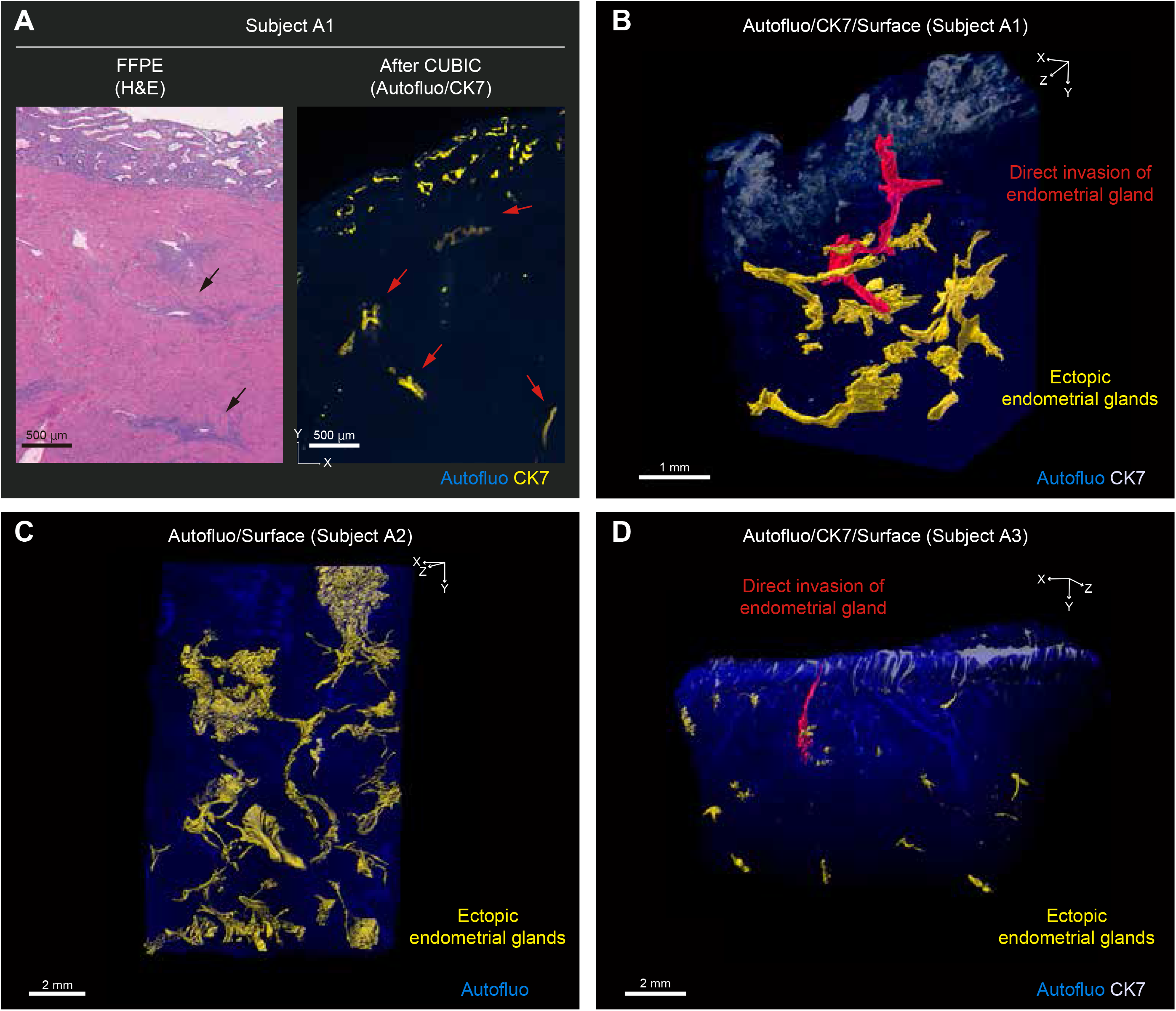
3D morphology of adenomyosis. (A) Left panel: Microscopic image of FFPE tissue after H&E staining of adenomyosis in the secretory phase (subject A1). Right panel: the reconstituted XY section (z = 10 μm) of the adenomyosis case after clearing by CUBIC. FFPE made by adjacent uterine tissue used for whole-mount 3D analysis. Black and red arrows indicate adenomyotic lesions. (B-D) 3D distribution of adenomyosis. (B) Subject A1. (C) Subject A2. (D) Subject A3. The subject A2 sample did not include the eutopic endometrium. Red object: 3D structures of the direct invasion of the endometrial gland into the myometrium. Yellow and red objects made by the Surface module in the Imaris software. After surface extraction, each structure was manually curated, and extra surface signals were eliminated. Images were obtained by LSF microscopy. Autofluorescence was measured by excitation at 488 nm. The CK7-expressing endometrial epithelial cells were measured by excitation at 532 nm lasers. FFPE, formalin-fixed paraffin-embedded; H&E, hematoxylin and eosin; Autofluo, autofluorescence; CK7, cytokeratin 7 See also Figure S6.

## Discussion

In this study, we succeeded in rendering a full-thickness 3D image of the human endometrium by using an updated CUBIC protocol and LSF microscopy. Our 3D imaging revealed characteristic morphological features of human endometrial glands, including the occluded glands and plexus of the basal glands, which were not sufficiently observed by 2D histology alone. On the other hand, these morphological features were detected regardless of age or menstrual cycle phase, suggesting that they were basic components of the normal human endometrium.

The 2D shape of the endometrial gland, as shown in Figure 6A, has been described from the early 1900s until now (Gray, 1918; Lessey and Young, 2019; Manconi et al., 2003; Padykula et al., 1984; Ross and Reith, 1985). However, our 3D image of endometrial glands suggests that this conventional 2D shape of the endometrial gland does not reflect the true morphology of endometrial glands. On the basis of our 3D observation, we can provide a new 2D shape of the human endometrial glands, which is the first in nearly one hundred years (Figure 6B). Specifically, we referred to the plexus as the ‘rhizome’ because of the similarity between the plexus and the rhizome in terms of not only their morphologies but also their functional features; for example, rhizomatous plants, such as grass, are able to regenerate from erosion (Yu et al., 2008).

**Figure 6.**
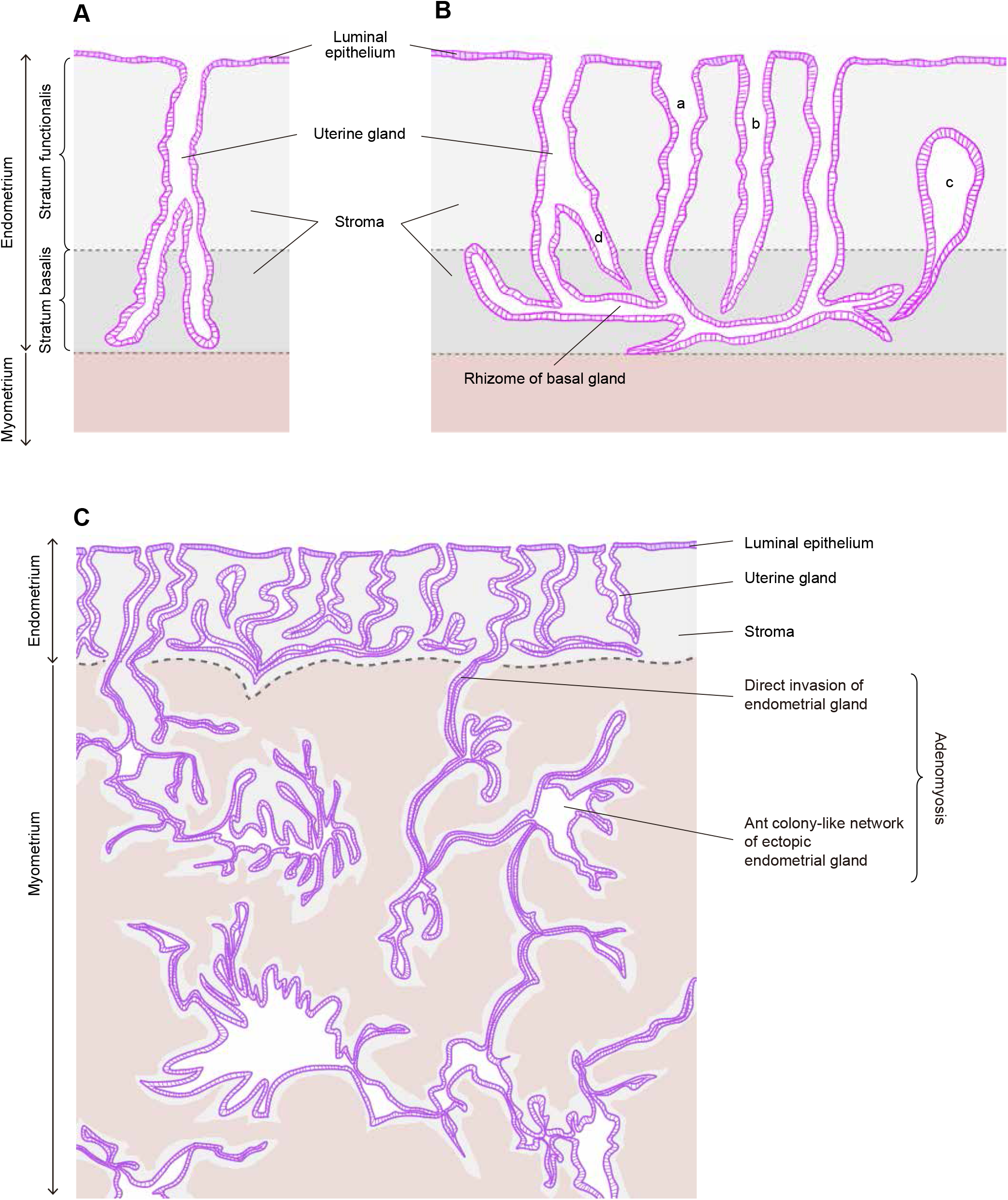
2D-shape image of the normal human endometrium and adenomyosis. (A) Conventional 2D-shape image of the endometrium (B) New 2D-shape image of the endometrium (C) 2D-shape image of adenomyosis including direct invasion of endometrial glands into the myometrium and an ant colony-like network of ectopic endometrial glands within the myometrium. a: Gland with rhizome, b: Gland with no rhizome, c: Occluded gland, d: Branch

In some studies, reconstituted 3D visualization of the partial endometrial structures was performed by using computerized two-dimensional binary images of serial sections or multiphoton excitation microscopy (Manconi et al., 2003; Manconi et al., 2001; Simbar et al., 2004). However, these studies had serious limitations in that the observable depth was less than 120 μm because of tissue transparency and could not detect the detailed 3D structure of the endometrial gland. Recently, several groups have developed tissue-clearing techniques (Chung et al., 2013; Erturk et al., 2012; Hama et al., 2011; Ke et al., 2013; Susaki et al., 2015; Tainaka et al., 2016). These techniques make a whole organ or sample transparent so that light could illuminate deep regions of the tissues. To date, only Arora *et al*. have applied tissue clearing and confocal imaging methods to only one human uterine tissue in addition to several mouse uterine tissues (Arora et al., 2016). However, they did not clarify any characteristic 3D morphologies of human endometrial glands. By using an updated CUBIC protocol (Tainaka et al., 2018), we successfully cleared human uterine tissue at a depth of several centimeters. The autofluorescence signal that remained slightly after the CUBIC protocol was useful to observe the details of anatomical structures in the tissue. Furthermore, we performed immunohistochemistry with a fluorescently labeled anti-CK7 antibody to extract the clear tubular structures of endometrial glands. As a result, we could make 3D surface rendering of endometrial glands by Imaris software (Bitplane).

Histologically, isolated cystically dilated glands are commonly encountered in normal endometrium. In this study, according to 3D visualization, we proved that the cystically dilated gland was the occluded gland, which was not continuous with the luminal epithelium. Because the occluded glands cannot discharge a secretion into the uterine cavity, a secretion accumulates and inspissates in the gland, leading to cystic dilatation of the gland. Cystically dilated glands are predominantly detected in the atrophic endometria of postmenopausal women and the disordered proliferative endometrium (Al-Hussaini et al., 2020). The irregular shapes and sizes of glands, including cystic dilatation without nuclear atypia, are characteristic of simple endometrial hyperplasia (Chandra et al., 2016). Endometrial hyperplasia frequently results from chronic estrogen stimulation unopposed by the counterbalancing effects of progesterone (Sanderson et al., 2017). Therefore, gland occlusion might be related to hormone imbalance and/or hormone sensitivity of the gland.

In the stratum basalis of the human endometrium, narrow and horizontally running glands are often detected by 2D histological observation (Garry et al., 2010). However, it has not been noted that the branches of glands form a complicated pattern in the stratum basalis. This is the first report to explicitly refer to the rhizome, which is the plexus morphology of basal glands proven by 3D observation. Although previous studies have shown the 3D structure of murine endometrial glands (Arora et al., 2016; Vue et al., 2018), the bottom of the murine endometrial gland forms a crypt but not a rhizome. This can be potentially explained by the existence of menstruation, which is the crucial difference between the human and murine endometrium. The human endometrium is a dynamic remodeling tissue that undergoes more than 400 cycles of regeneration, differentiation, shedding, and rapid healing during a woman’s reproductive years (McLennan and Rydell, 1965). Because the stratum functionalis is shed during menses, the endometrium is believed to regrow and regenerate from endometrial progenitor/stem cells residing in the stratum basalis (Kyo et al., 2011; Padykula, 1991; Padykula et al., 1984; Prianishnikov, 1978). Although it has been established that intestinal stem cells exist in the bottom of the crypt (Sangiorgi and Capecchi, 2008), there are still uncertainties about the localization of endometrial progenitor/stem cells (Santamaria et al., 2018). Therefore, it is necessary to take into consideration that the human endometrium has a rhizome in its stratum basalis in future endometrial stem/progenitor cell studies. Furthermore, rhizomatous plants that have stems running underground horizontally are well known for their difficulty in terms of eradication because they can regenerate from a piece of rhizome left behind in the soil after natural or artificial erosion (Sásik and Elias, 2006; Yu et al., 2008). If endometrial progenitor/stem cells are located in the stratum basalis, the rhizome in the human endometrium will have a functional advantage over the crypt in terms of the conservation of progenitor/stem cells and regeneration.

Some studies have indicated that the human endometrial epithelial glands are monoclonal in origin (Chan et al., 2004; Gargett et al., 2016; Kyo et al., 2011; Tanaka et al., 2003), implying that they arise from a single progenitor/stem cell. Our group has recently reported diversification of cancer-associated mutations in histologically normal endometrial glands (Suda et al., 2018). In our previous study, we sequenced 109 single endometrial glands isolated from the stroma using collagenase and found that two of them shared the same *PIK3CA* mutation (p.K111N) and the same *PPP2R1A* mutation (p.S256Y), with high mutant allele frequencies. It was suggested that both glands descended from a single common ancestral cell. If an endometrial gland has a monoclonal composition including rhizome, it follows that some glands sharing the rhizome have a common origin and that the endometrium is an aggregate of small clonal segments. The genomic alterations of the progenitor/stem cell may transmit to several glands through a rhizome. Indeed, the most recent study of the whole-genome sequencing of the normal human endometrial epithelium showed that six microdissected glands that were isolated from one section shared over 100 variants; therefore, they were regarded as the same clade (Moore et al., 2020). The authors argued that clonal evolution of phylogenetically related glands entailed the capture and colonization of extensive zones of the endometrial lining (Moore et al., 2020). Interestingly, their phylogenetically related glands were located at the bottom of the stratum basalis and interspersed horizontally, suggesting that they form rhizomes in 3D observations. It is possible that the rhizome of the endometrium is a crucial element for understanding the genetic features of the endometrium.

In this study, we also succeeded in observing the 3D morphology of adenomyosis by our method. Adenomyosis is defined as the presence of ectopic endometrial glands and stroma surrounded by hyperplastic smooth muscle within the myometrium (Vannuccini et al., 2017). There are several hypotheses about the etiology of adenomyosis, such as endometrial invasion, endometriotic invasion, and de novo metaplasia (Kishi et al., 2012). Among them, it has been generally accepted that uterine adenomyosis results from direct invasion of the endometrium into the myometrium. This pathologic condition was advocated based on an observation of 2D serial sections (Benagiano and Brosens, 2006). In this study, we successfully proved the direct invasion of the endometrium into the myometrium in reconstituted 3D images of CK-7-stained adenomyosis. We also depicted how adenomyosis expands lesions within the myometrium. Our 3D images showed that the ectopic endometrial glands had lengthened thin branches and expanded lesions with an ant colony appearance in patients not undergoing hormone therapy. Based on our 3D images, we could provided a new 2D shape of adenomyosis shown in Figure 6C.

A recent study involving genomic analysis of adenomyosis indicated high variant allele frequencies of *KRAS* hotspot mutations in the microdissected epithelial cells of adenomyotic tissue, which suggested that adenomyosis may arise from the ectopic proliferation of mutated epithelial cell clones (Inoue et al., 2019). Furthermore, they found identical *KRAS* mutations in adenomyotic and histologically normal endometrium adjacent to adenomyotic lesions and argued that “*KRAS*-mutated adenomyotic clones originate from normal endometrium” (Inoue et al., 2019). Our findings of the 3D morphology of adenomyosis support this hypothesis from a histological perspective. A combination of genomic and 3D analysis is required to elucidate the etiology of adenomyosis.

This study has some limitations that must be addressed. First, we could not perform whole-uterine clearing or whole-uterine sampling because our samples were obtained clinically from patients who underwent hysterectomy. Structural mapping of the whole human uterine endometrium improves the understanding of the anatomical and histological features of the human uterus. Second, our samples did not include young women under 29 years old since young women seldom undergo hysterectomy. Third, we revealed structural details of the human endometrial glands by anti-CK7 antibody labeling, but no cellular or molecular analysis was performed. Because in vivo genetic labeling and fluorescent dye tracing are not applicable to human studies (Susaki and Ueda, 2016; Zhao et al., 2020), cellular and molecular analysis of human organs requires a postsampling staining method based only on diffusion penetration using fluorescently labeled antibodies in updated CUBIC protocols (Tainaka et al., 2018). Since antigenicity is dependent on histological preparation conditions (e.g., fixation and clearing), the detailed histological preparation conditions should be described for each working antibody (Nojima et al., 2017). Further search of the antibody suitable for the 3D staining of centimeter-sized human tissues will be necessary in future cellular and molecular analyses. Finally, this study did not connect the 3D morphology of the glands with genomic data. Spatial genomics and transcriptomics are hot topics in omics research (Stahl et al., 2016). Construction of the 3D genomics and/or transcriptomics pipeline is expected.

In conclusion, we successfully obtained 3D full-thickness images of the human endometrium using an updated CUBIC protocol. This stereoscopic imaging made it possible to analyze free-angle images of cross-sections. With this procedure, we can visualize the 3D morphology of the glands and create the new concept of 2D-shape images of the human endometrium. Furthermore, using our protocol, we revealed the 3D pathologic morphology of adenomyotic lesions. From these findings, we conclude that this procedure is a useful tool to analyze the human endometrium and endometrium-related diseases from a new perspective. The 3D representation of the human endometrium will lead to a better understanding of the human endometrium in various fields, including histology, pathology, pathophysiology and oncology.

## Acknowledgments

This work was supported in part by the Japan Society for the Promotion of Science (JSPS) KAKENHI grant number JP17H04336 (Grant-in-Aid for Scientific Research B for TE) JP19K09822 (Grant-in-Aid for Scientific Research C for KY). We are grateful to Anna Ishida and Kenji Ohyachi for their technical assistance.

## Author Contributions

Conceptualization, M.Y., K.Y., and T.E.; Methodology, M.Y., K.Y., and K.T.; Investigation, M.Y., K.Y., and K.T.; Validation, T.M.; Writing – Original Draft, M.Y. and K. Y.; Writing – Review & Editing, K.T., and T.E.; Funding Acquisition, K. Y., and T.E.; Resources, K. Suda, K. Sugino, N.Y., R.T., T.I.; Supervision, K.Y., H.N., K.T., and T.E.

## Lead contact

Further information and requests for resources and reagents should be directed to and will be fulfilled by the Lead Contact, Kosuke Yoshihara.

## Materials Availability

The data on 3D histology of human uterine endometrium and endometrium-related diseases are freely available from the Lead contact and shared at TRUE (Three-dimensional Representation of human Uterine Endometrium) that is a database web site (https://true.med.niigata-u.ac.jp/).

## Methods

### Human sample collection and histological examination

This study was approved by the institutional ethics review boards of Niigata University. We recruited study participants at the Niigata University Medical and Dental Hospital between August 2018 and September 2019. All subjects provided written informed consent for the collection of samples and analyses.

We collected 14 uterine endometrial samples from 12 patients (aged 30-49 years) with a nonendometrial gynecological disease who underwent hysterectomy. Adenomyosis samples were collected from three patients (42-45 years old). Each sample was divided into two blocks: one was used for whole-mount 3D analysis, and the other was used for histological examination. The fresh human tissues in the latter block were fixed in neutral formalin and embedded in paraffin. They were then used for staining with hematoxylin and eosin (H&E) and for a series of immunostainings. Histological diagnoses, including menstrual cycle phase, were reviewed by an experienced gynecologic pathologist (T.M.).

### CUBIC protocol for whole-mount 3D staining with CK7

The updated CUBIC protocols were previously described (Tainaka et al., 2018). We applied CUBIC protocol IV, which is suitable for clearing human brain tissue. Human endometrium blocks (5-14.4 mm× 4.7-11.9 mm ×3.9-13.1 mm) and adenomyosis blocks (7.3-17.9 mm × 7.9-18.6 mm × 5.6-16.9 mm) were stored in formalin until use. The tissue blocks were washed with PBS for 6 hours before clearing. Then, the tissue blocks were immersed in CUBIC-L [T3740 (mixture of 10 wt% *N*-butyliethanolamine and 10 wt% Triton X-100), Tokyo Chemical Industry] with shaking at 45 □ for 6-8 days. During delipidation, the CUBIC-L was refreshed once. After the samples were washed with PBS for several hours, the tissue blocks were placed into 1-3 ml of immunostaining buffer (mixture of PBS, 0.5% Triton X-100, 0.25% casein, and 0.01% NaN_3_) containing 1:100 diluted Alexa647 or 555-conjugated cytokeratin (CK) 7 antibody (ab192077 or ab203434, Abcam) for 10-14 days at room temperature with gentle shaking. After the samples were washed with PBS for several hours, the samples were subjected to postfixation by 1% PFA in 0.1 M PB at room temperature for 5 hours with gentle shaking. The tissue samples were immersed in 1:1 diluted CUBIC-R+ [T3741 (mixture of 45 wt% 2,3-dimethyl-1-phenyl-5-pryrazolone, 30 wt% nicotinamide and 5 wt% *N*-butyldiethanolamine), Tokyo Chemical Industry] with gentle shaking at room temperature for 1 day. The tissue samples were then immersed in CUBIC-R+ with gentle shaking at room temperature for 1-2 days.

### Microscopy

Macroscopic whole-mount images were acquired with a LSF microscope (MVX10-LS, Olympus). Images were captured using a 0.63 × objective lens [numerical aperture (NA) = 0.15, working distance = 87 mm] with digital zoom from 1 × to 3.2 ×. The LSF microscope was equipped with lasers emitting at 488 nm, 532 nm, and 637 nm. When the stage was moved to the axial direction, the detection objective lens was synchronically moved to the axial direction to avoid defocusing. Alexa 555 or 647 signals of CK7-expressing endometrial epithelial cells were measured by excitation at 532 nm or 637 nm lasers. Autofluorescence was measured by excitation at 488 nm or 532 nm (if CK7 was expressed at 637 nm).

### Image analysis

All raw image data were collected in a lossless 16-bit TIFF format. All fluorescence images of CK7 were obtained by subtracting the background and unsharp mask using Fiji software. 3D-rendered images were visualized, captured and analyzed with Imaris software (version 9.3.1 and 9.5.1, Bitplane). Image analysis by Imaris software was previously described (Arora et al., 2016; Tainaka et al., 2014), and we modified it. TIFF files were imported into the Surpass mode of Imaris. The reconstituted 3D images were cropped to a region of interest using the 3D Crop function. Using the channel arithmetic function, the CK7 signal was removed from the autofluorescence signal to create a channel with only endometrial epithelium and gland signals. The extracted glands were then 3D-reconstracted by the Surface module. After surface extraction, each structure was manually curated, and extra surface signals were eliminated. When 3D surface objects were made in the Imaris software, disconnected components could be selected individually for assignment of a pseudocolor and for separation into new channels. Thus, each occluded gland, namely, glands with rhizome or an adenomyotic lesion, were pseudocolored individually. The snapshot and animation function were used to capture images and videos, respectively.

### 3D-layer distribution

To describe the horizontal morphology of uterine glands from the basalis to luminal epithelium, Imaris XT software was adapted for our use. This module is a multifunctional two-way interface from Imaris to classic programming languages such as MATLAB, Python or Java that enables users to rapidly develop and integrate custom algorithms that are specific and tailored to scientific applications where generic image processing would fail. We chose the distance transformation (DT) tool from Imaris XT tools for layer distribution analysis. First, a 3D tissue image was cropped in the XZ plane to 2.5 × 2.5 mm, and a 3D bottom of the endometrium was created using the manually Surface module. The borderline between the endometrium and myometrium was traced every 25 slices or less in the XY plane. After the 3D bottom surface was generated, the DT tool was selected from the same surface tools as the outside surface object mode, and then, a new DT channel was created. Second, 3D layers of the endometrium were created using the surface module with the newly created DT signal. The threshold was set by a width of 150 μm. Thus, a new 3D-layer surface was created at the same distance from the bottom layer and thickness of 150 μm. Third, the mask channel module was applied to each layer with the CK7 signal. Each new layer of the uterine glands was separated and pseudocolored. Finally, 3D morphological images of the uterine glands of each layer were reconstituted using the Surface module with the newly created CK7 signal of each layer.

## Supplemental Information

**Figure S1**

The reconstituted XY-section images (z = 1.7 μm) of human uterine tissue (subject E7) stained with Alexa Fluor 647-conjugated anti-CK7 antibody with clearing by CUBIC. Images were obtained by LSF microscopy with single-cell resolution. Autofluorescence was measured by excitation at 532 nm. The CK7-expressing endometrial epithelial cells were measured by excitation at 637 nm.

Autofluo, autofluorescence; CK7, cytokeratin 7

**Figure S2**

The occluded glands of the proliferative phase (subjects E1 to 4) and secretory phase (subjects E5-1 to 7, 9 and 10). The reconstituted 3D morphologies of the occulated glands were pseudocolored independently by Imaris software. Images were obtained by LSF microscopy. Autofluorescence was measured by excitation at 523 nm (subject E7) and 488 nm (all other subjects). The CK7-expressing endometrial epithelial cells were measured by excitation at 647 nm (subject E7) and 523 nm (all other subjects). Autofluo, autofluorescence; CK7, cytokeratin 7

**Figure S3**

The plexus structure of the glands in the proliferative phase (subjects E1 to 4) and secretory phase (subjects E5-1 to 7, 9 and 10). The reconstituted XY-section images (z = 100 μm). Red arrows indicate the branches. Images were obtained by LSF microscopy. Autofluorescence was measured by excitation at 523 nm (subject E7) and 488 nm (all other subjects). The CK7-expressing endometrial epithelial cells were measured by excitation at 647 nm (subject E7) and 523 nm (all other subjects).

Autofluo, autofluorescence; CK7, cytokeratin 7

**Figure S4**

Making of 3D-bottom and layer surface of human endometrial glands (subject E1).

(A, B) The reconstituted 3D bottom surface of the endometrium. The borderline between the endometrium and myometrium of the XY section was manually traced by the Surface module in the Imaris software.

(C, D) Making of the 3D-layer surface of the distance transformation channel at the same distance from the bottom layer and with a thickness of 150 μm.

Layer 1 (magenta): 1-150 μm; layer 2 (green): 151-300 μm; layer 3 (light blue): 501-650 μm; layer 4 (orange): 1001-1150 μm; and layer 5 (yellow): 1501-1650 μm. (E, F) The reconstituted 3D-layer distribution of endometrial glands.

Images were obtained by LSF microscopy. Autofluorescence was measured by excitation at 488 nm. The CK7-expressing endometrial epithelial cells were measured by excitation at 532 nm lasers.

Autofluo, autofluorescence; CK7, cytokeratin 7

**Figure S5**

3D-layer distribution of endometrial glands in menstruation cases (subjects E11-2 and 12). (A) Subject E11-2 (menstrual cycle day 2). (B) Subject E12 (menstrual cycle day 4). Left panels: Microscopic image of FFPE tissue after H&E staining. Middle panels: The reconstituted XY section (E11-2, z = 4.3 μm; E12, z = 5.6 μm) after clearing by CUBIC. Each layer was pseudocolored by Imaris software. Layer 1 (magenta): 1-150 μm; layer 2 (green): 151-300 μm; and layer 3 (light blue): 501-650 μm. Right panels: The XZ plane view (y = 150 μm) of layers made by the Surface module in the Imaris software. After surface extraction, each structure was manually curated, and extra surface signals were eliminated.

Images were obtained by LSF microscopy. Autofluorescence was measured by excitation at 488 nm. The CK7-expressing endometrial epithelial cells were measured by excitation at 532 nm.

Autofluo, autofluorescence; CK7, cytokeratin 7

**Figure S6.**

3D morphology of adenomyosis. (A) Subject A2. (B) Subject A3. The subject A2 sample did not include the eutopic endometrium. Left panels: Microscopic image of FFPE tissue after H&E staining. Middle panels: The reconstituted XY section (A2, z = 9.2 μm; A3, z = 15.7 μm) of the adenomyosis case after clearing by CUBIC. Black and red arrows indicate adenomyotic lesions. Right panels: 3D distribution of adenomyosis. Images were obtained by LSF microscopy. Autofluorescence was measured by excitation at 488 nm. The CK7-expressing endometrial epithelial cells were measured by excitation at 532 nm.

FFPE, formalin-fixed paraffin-embedded; H&E, hematoxylin and eosin; Autofluo, autofluorescence; CK7, cytokeratin 7

**Video S1**

Reconstituted 3D image of proliferative-phase full-thickness human endometrial tissue (subject E2) stained with CK7 (yellow) and showing autofluorescence (blue). Z-stack (XY plane view): 100 μm.

**Video S2**

Reconstituted 3D image of the plexus of the endometrial glands (subject E8) stained with CK7 (gray) and showing autofluorescence (blue). The glands sharing the plexus were pseudocolored individually (green or light blue). Z-stack (XY plane view): 6.61 μm.

**Video S3**

3D-layer distribution of endometrial glands in menstruation cases (subject E11-1).

Yellow: CK7, Blue: autofluorescence. Z-stack (XY plane view): 52.8 μm.

**Video S4**

Reconstituted 3D image of adenomyotic lesion stained with CK7 (gray) and showing autofluorescence (blue). The adenomyotic lesions were pseudocolored individually (red: direct invasion of the endometrial gland into the myometrium, yellow: ectopic endometrial glands). Z-stack (XY plane view): 10 μm.

